# Reconstructing Langevin systems from high and low-resolution time series using Euler and Hermite reconstructions

**DOI:** 10.1101/2024.05.30.596585

**Authors:** Babak M. S. Arani, Stephen R. Carpenter, Egbert H. van Nes

## Abstract

The ecological literature often features phenomenological dynamic models lacking robust validation against observational data. Reverse engineering is an alternative approach, where time series data are utilized to infer or fit a stochastic differential equation. This process, known as system reconstruction, presents significant challenges. This paper addresses the estimation of the (often) non-linear deterministic and stochastic parts of Langevin models, one of the simplest yet commonly used stochastic differential equations. We introduce a Maximum Likelihood Estimation (MLE) inference method, termed Euler reconstruction, tailored for time series data with medium to high resolution. However, the Euler approach is not reliable for low-resolution data. To fill the gap for sparsely sampled data, we present an MLE inference method pioneered by Aït-Sahalia, that we term Hermite reconstruction. We employ a powerful modeling framework utilizing splines to detect inherent nonlinearities in the unknown data-generating system to achieve high accuracy with minimal computational burden. Applying Euler and Hermite reconstructions to a range of simulated, ecological, and climate datasets, we demonstrate their efficacy and versatility. We provide a user-friendly tutorial and a MATLAB package called the ‘MATLAB reconstruction package’.

## Introduction

It has long been a matter of debate whether ecosystems can have alternative stable states, how to measure their resilience and whether they can recover from perturbations. These are fundamental ecological questions which have mainly been discussed using theoretical models (Connell & Sousa 1983). Tackling these questions necessitates a reverse engineering approach, where we reconstruct the system based on dynamic data (Siegert & Friedrich 2001; Rinn *et al*. 2016). This approach involves elucidating the unknown data-generating system using dynamic data. Subsequently, the resilience of the best-fitting model can be studied (Bolker *et al*. 2013; Hilborn & Mangel 2013). When appropriate mechanistic dynamical equations are known, parameters of the equations can be inferred from time series data.

Ecologists are often confronted with a situation where the nonlinear stochastic model that generated the data is unknown and must be inferred from data. If data are measured frequently throughout the entire range of the state variables, then a nonlinear stochastic model can be reconstructed by nonparametric methods (Bandi & Phillips 2003; Rinn *et al*. 2016). The reconstructed model can then be analyzed to map the stability landscape, locate stable or unstable equilibria, and calculate stochastic indicators of resilience such as mean exit time or median survival time (Arani *et al*. 2021). While nonparametric reconstruction methods are powerful, they have several limitations. They require high-resolution data and involve arbitrary choices, such as estimating the conditional mean, conditional variance, and higher conditional moments using kernels with bandwidths that control smoothness. Estimating the bandwidth is challenging, and the results may be sensitive to its choice. Additionally, these methods extrapolate moments to zero for a specified first few numbers of lags, defined as integer multiples of the time step in the data (Siegert & Friedrich 2001; Bandi & Phillips 2003; Rinn *et al*. 2016). The choice of the number of lags significantly affects the results, particularly when data resolution is low. Furthermore, these methods demand substantial amounts of data for reliable estimation (Siegert et al., 1998; Rinn et al., 2016). An alternative approach involves model-fitting techniques that maximize a likelihood function. This approach includes both the classical Euler scheme and a novel methodology pioneered by Aït-Sahalia (Aït-Sahalia 2002). It offers an attractive method that requires fewer data for reliable parameter estimation.

This paper presents a maximum likelihood estimation (MLE) approach for reconstructing the following one-dimensional Langevin models from time series data

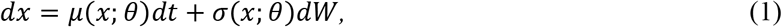

where *μ*(*x*;*θ*) denotes the deterministic component of the system, referred to as the ‘drift function’ and *σ*(*x*;*θ*) represents the stochastic component, known as the ‘diffusion function’. *θ* represents the parameter vector to be estimated via MLE and *W* refers to a Wiener process, where the noise source *dW*, known as Brownian noise, is Gaussian and white (uncorrelated). The diffusion function *σ*(*x*;*θ*) weighs the impact of perturbations, reflecting the intensity of perturbations at state *x*. The noise in a diffusion model (1*11*) is additive if the diffusion function does not depend on the state *x* (i.e., it is constant in terms of state), otherwise, it is called multiplicative. The inference technique employed here is parametric meaning that a parametric form, with a vector of parameters *θ*, for the drift *μ*(*x*;*θ*) and diffusion *σ*(*x*;*θ*) functions should be predefined and then the parameters are estimated using time series data through MLE.

In this study, we apply the classical inference of Langevin models using Euler methodology which we call ‘Euler reconstruction’. This method requires data of medium or high resolution, posing a significant challenge in fields such as ecology and climate science. To address this issue, we present an MLE scheme based on the groundbreaking work of Aït-Sahalia (Aït-Sahalia 2002) for univariate sparsely sampled data. The idea of Aït-Sahalia involves constructing a convergent series expansion of the conditional densities of (1) using Hermite polynomials, so we call this procedure ‘Hermite reconstruction’. To our knowledge this paper is the first to provide an open-access MATLAB package and to demonstrate successful applications to ecosystem and climate data. One of our key techniques for addressing the underlying MLE problem is the utilization of a wide variety of spline models as representation of the unknown drift and diffusion functions. Splines are flexible non-linear structures in terms of the state variable *x* and are linear functions of the parameter vector *θ*, significantly enhancing the accuracy and speed of calculations (as demonstrated in various examples in our user-friendly tutorial). Additionally, splines serve as valuable tools in cases where selecting an appropriate model is uncertain, which is often encountered. To distinguish this modeling technique from typical ‘parametric modeling’ we term it ‘spline modeling’.

The remainder of the paper presents the steps of the analysis, demonstrates reconstruction of diffusion models for several simulated, ecological, and climate data and shows that the method succeeds even for rarified datasets. Supplementary materials include a mathematical appendix, and a tutorial to illustrating a broader range of the capabilities of the Euler and Hermite reconstructions using both parametric and spline models across different data types. These data types encompass typical time series data (i.e., single datasets), replicate time series (multiple datasets belonging to the same data-generating system), big datasets, datasets with missing values, or possible combinations thereof.

### Steps of the Method

#### Step 1. Prepare the data: data standardization

In some real datasets, the scale of the data can be very large. In such cases, data standardization, achieved by computing z-scores (subtracting the mean and dividing by the standard deviation), makes it easier to solve the MLE. This is because standardization helps to narrow down large search spaces, making them more manageable. Additionally, standardization brings the data to a common scale, centered at 0 with small dispersion. This, in turn, makes it convenient to define a small region of parameter space for the MLE algorithm to search within (See Step4 for more details on this). For further illustration on data standardization, refer to section 8 of tutorial and Appendix G if you are interested in technical details

#### Step 2. Check the data requirements

There are three data requirements for a Langevin model in (*1*) which should be checked prior to performing the reconstruction (for a detailed discussion see section 6 of the tutorial). Firstly, data should be stationary, meaning that its statistical properties remain unchanged throughout the study period. If this assumption is violated, the data should be divided into smaller (possibly overlapping) periods where stationarity is assured. Reconstruction can then be carried out separately for each period, with the final system reconstruction obtained by interpolating the results. Secondly, data should be Markovian meaning that the future state of data, given the present state, should be independent of the entire past history of states. The smallest time scale at which Markovicity holds is called ‘Markov-Einstein’ (ME) time scale (Friedrich *et al*. 2011). Reconstruction can, then, be performed on a rarified sample of data whose resolution matches the ME time scale or on samples with lower resolutions. Thirdly, it is essential to check the resolution of the rarified sample regarded now as our dataset to be analyzed. This is the subject of next step.

#### Step 3. Check the resolution of data: how high should the data resolution be?

How high should the data resolution be in order to be able to reconstruct the data-generating system? This is a question we should investigate prior to performing any reconstruction procedure. The answer depends on the ‘speed’ or ‘time scale’ of the yet unknown system relative to sampling frequency of the data. To investigate the time scale of data the autocorrelation of data should be examined and a quantity known as ‘*relaxation time*’ *τ_R_* should be estimated. For univariate data, we can roughly estimate relaxation time directly from data by fitting the exponential exp(−*ct*) to the initial lags of the data autocorrelation function, obtaining the estimate *ĉ*, and setting the relaxation time *τ_R_* = 1/*ĉ* (refer to Appendix A for the details). Assuming Δ to be the data sampling time we consider the following three regimes.

##### The first regime: Δ being much smaller than τ_R_

This is the desired regime and we can safely apply simple reconstruction schemes like Euler scheme or Langevin approach. Unfortunately, many real datasets are not sampled frequently enough to fall into this regime. Furthermore, some high-resolution data do not adhere to the Markov property, which is another data requirement (see Step 3). However, a rarified sample of such data may be Markovian but this comes at the expense of reduced resolution.

##### The second regime: Δ is approximately the same order of magnitude as τ_R_

In theory, this represents the minimum resolution at which we can recover the data-generating system. In such cases, a more accurate reconstruction procedure is necessary compared to the Euler scheme. Here, Hermite reconstruction becomes essential. Refer to section 11.2 of the tutorial, which showcases a dataset accurately reconstructed with high precision using Hermite reconstruction at a resolution matching *τ_R_* .

##### The third regime: Δ being bigger than τ_R_

In this regime consecutive measurements are almost independent. Therefore, the true dynamics is not reflected in such datasets and any reconstruction procedure is expected to fail.

To perform a successful reconstruction, the resolution of the dataset should fall between the first two regimes. In Section 6.3 of the tutorial, we provide a convention for categorizing data resolution into three categories: low, medium, and high. Euler reconstruction is suitable for datasets categorized as medium or high resolution. However, for datasets categorized as low resolution, we employ Hermite reconstruction.

#### Step 4. Guidelines for implementing parametric and spline modeling

In parametric modeling, parametric forms for the drift and diffusion functions in the Langevin model (1*11*) should be specified prior to embarking on MLE. For instance, in the Ornstein-Uhlenbeck model the parametric forms for the drift and diffusion functions are *μ*(*x*) = −*μx* and *σ*(*x*) = *σ*, respectively where *θ* = [*μ, σ*] is the vector of parameters. Similarly, in the grazing model of May (May 1977) 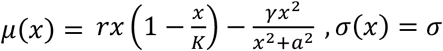, *σ*(*x*) = *σ* where *x* represents the biomass of a plant population, *θ* = [*r, K, γ, a, σ*] is the vector of parameters in which, *r* is the growth rate, *K* is the carrying capacity, *γ* is the maximum grazing rate, *a* is the efficiency of the grazer, and *σ* is the intensity of environmental perturbations which is assumed to be constant (so, this is an additive model). In spline modeling, no model should be specified. Instead, a rather sparse mesh across the range of data, called ‘knot sequence’, is defined. Knots are the x-coordinate of the points where spline interpolation goes through. The values of the spline function at the knots are the model parameters (see Figures 1-5). Splines, therefore, offer flexibility in capturing unknown nonlinearities in drift and diffusion functions, making them particularly attractive when a suitable model is uncertain. Note that spline modeling is also parametric but to distinguish its rather distinct features from ‘typical’ parametric models we call it that way.

**Figure 1.**
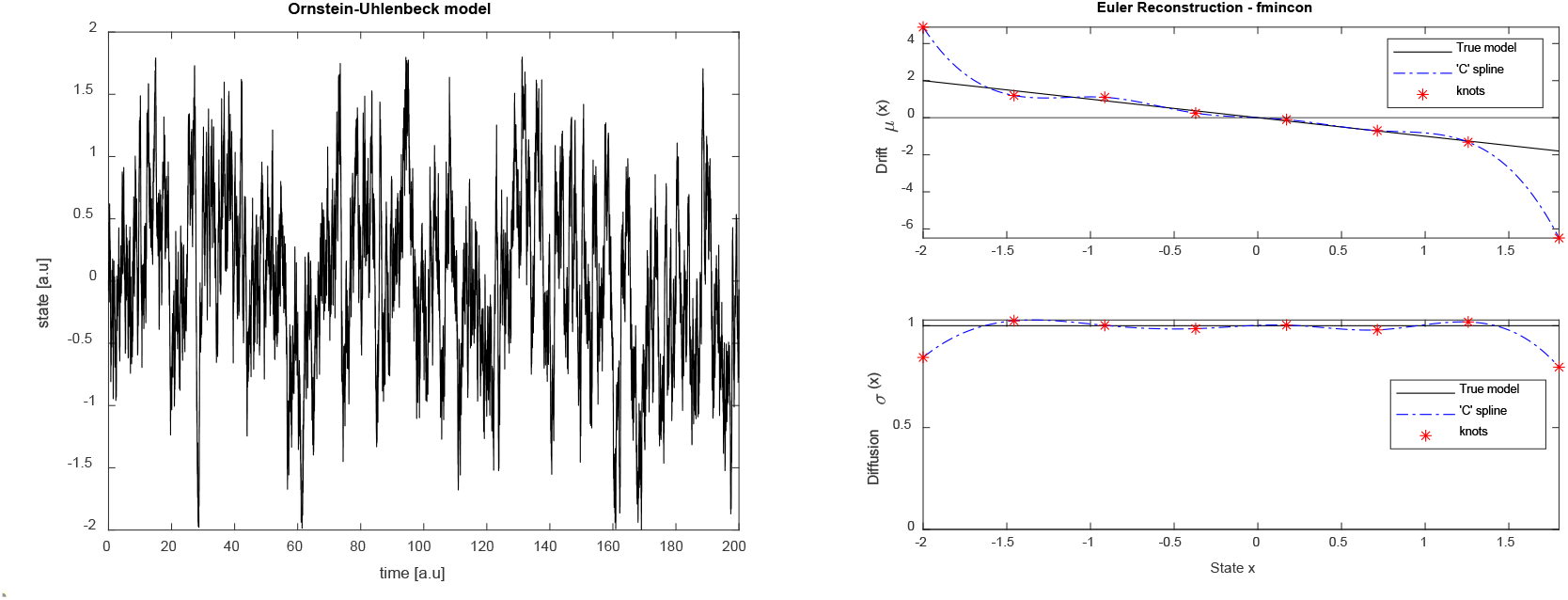
Illustration of spline reconstruction for a high-resolution dataset generated from a linear model. Data are illustrated in the left panel where 20000 data points, with sampling time Δ = 0.01, are generated from the OU model. The right panel illustrates estimated cubic spline models (dot-dashed blue curves) for both the drift and diffusion functions using 8 regularly spaced knots over the state space alongside the true drift and diffusion functions (black curves). a.u means arbitrary unit.

After selecting a model (parametric or spline), a vector of lower bounds and a vector of upper bounds for the parameters should be specified for the optimization (MLE) to search within and find the optimal parameter values. All parameters associated with the diffusion function should be bounded in a way that ensures the diffusion function remains positive. For parametric models, the physics of the problem can provide insights to determine proper vectors of lower and upper bounds. For example, in the case of the May model, we know that all model parameters should be positive as they represent ecological quantities. For spline models selecting vectors of lower and upper bounds are convenient. Since datasets we are dealing with are stationary then for the drift parameters, we can set all lower bounds to be -L and all upper bounds to be L, where L>0. Similarly, for the diffusion parameters, we can set all lower bounds to 0 and all upper bounds to L.

If a parametric modeling is preferred and it is uncertain to choose a proper model then it is advisable to select drift and diffusion models which contain constant, linear, quadratic, and higher order terms due to Taylor series expansion. An additive version of such a modeling is as follow

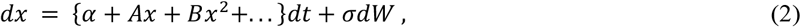

The parameter *A* is particularly insightful as it is the sole determinant of stability of the deterministic part of (2). The model form in (2) is not directly suited for fitting to data. Instead, data should be standardized first, and then the model (2) could be applied. Since, this model is linear in parameters, it is convenient to transform the estimated parameters back into their original scales by replacing the state variable *x* with its standardization (*x* − *m*)/*s* and multiplying the right-hand side by *s* where *m* and *s* are the mean and standard deviation of the data, respectively.

#### Step 5. Euler reconstruction

If the data resolution falls within either the high-resolution or medium-resolution category, then Euler reconstruction is applicable.

#### Step 6. Hermite reconstruction

If the data resolution falls in the category of low resolution, Hermite reconstruction should be employed. Here, we briefly outline the approach, but for detailed mathematical explanation, refer to Appendices F, G, H and I. For a friendly explanation refer to section 11 of the tutorial, which includes numerous case studies. In our MATLAB package, we have implemented the Hermite reconstruction based on a refinement by (Bakshi & Ju 2005) to a methodology developed by (Aït-Sahalia 2002).

Our approach involves a two-phase algorithm. In the first phase Euler reconstruction is performed, resulting in a first guess of the parameter vector called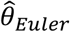. In the second phase, the algorithm improves this first guess using Hermite reconstruction. Hermite reconstruction requires two key inputs: J and K. J represents the number of ‘spatial’ terms in the Hermite expansion of the likelihood function using Hermite polynomials, while K represents the number of ‘temporal’ terms in the Taylor expansion, in Δ, of Hermite coefficients. High J and K increase estimation accuracy at the cost of higher computation time. Typically, a small J suffices, and in all case studies in the tutorial, we have used J=3. However, as data resolution decreases, a bigger K is necessary to enhance estimation accuracy. Based on our experience, for data with low-to-medium resolution a value of *K* ≤ 6 suffices while for extremely low-resolution data values of 6 < *K* ≤ 12 are needed. Consequently, reconstructing datasets with lower resolution becomes computationally more intensive. The primary challenge lies in dealing with an optimization problem with a partially-defined objective function in the second phase. As data resolution diminishes, the algorithm should search within smaller regions in the parameter space. To address this challenge, the algorithm first identifies a set of parameters, we call ‘legitimate points’ (LP), where the Hermite objective function has defined values.

Subsequently, the algorithm utilizes these LPs and searches in the vicinity of the Euler estimation 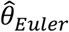 to get an improved vector of parameters 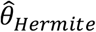. Refer to Appendix G For details of the optimization process.

Our package not only supports commonly used cubic splines but also offers a range of other spline types. The use of ‘quadratic’ splines is particularly significant for achieving considerably faster speeds compared to cubic splines in Hermite reconstruction. Finally, while both parametric and spline models can be used, spline modeling is generally more convenient, faster, and leads to greater accuracies.

### Examples with high-resolution data

#### Example 1

Here, we reconstruct a high-resolution but rather small dataset with 2000 data points (see Figure 1, left panel) generated from the OU model with drift function *μ*(*x*) = −*μx*, diffusion function *σ*(*x*) = *σ*, parameter values *μ* = *σ* = 1, and sampling time Δ= 0.01. We obtain the parameter estimation 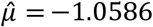 and 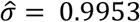 for this linear parametric model. We also perform a spline reconstruction to this dataset using 8 equidistance knots (see Figure 1, right panel).

#### Example 2

In this example we consider reconstructing a dataset (refer to Figure 2, left panel) simulated from the overgrazed model of May with drift function 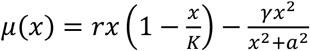 and diffusion function *σ*(*x*) = *σ* (in section ‘Step4’ the meaning of the parameters of May model were described). This dataset contains 10^5^ data points with sampling time Δ= 0.01. The parameter values are *r* = 1.01, *K* = 10, *γ* = 2.75, *a* = 1.6, *σ* = 0.4. Under these parameters the deterministic (i.e., without noise) model of May exhibits over-grazed and under-grazed alternative vegetation states (Figure 2, top right panel). Furthermore, the May model is nonlinear in terms of parameters *K* and *a* as well as the state variable *x*. Therefore, unlike the dataset in Example1, a longer dataset is needed here in order to be able to reconstruct transitions and time scales of shifts between alternative basins of attraction. Nonetheless, we could estimate parameters with a rather good accuracy as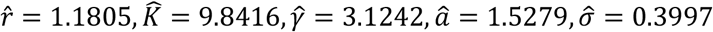. Figure 2, right panel illustrates a cubic spline model fitted to this dataset.

**Figure 2.**
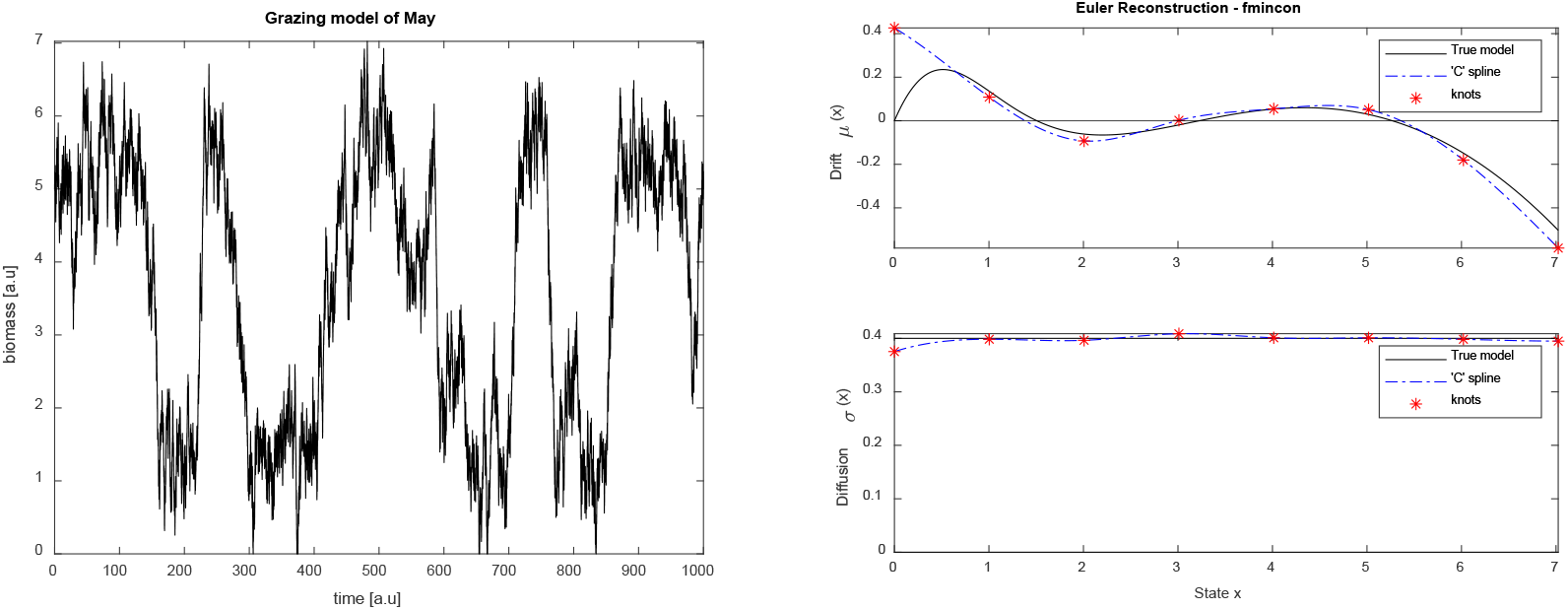
Illustration of spline modeling for a high-resolution dataset generated from a nonlinear model. Data are illustrated in the left panel where 10^5^ data points, with sampling time Δ = 0.01, are generated from the overgrazed model of May. The right panel illustrates estimated cubic spline models (dot-dashed blue curves) for both the drift and diffusion functions using 8 regularly spaced knots over the state space alongside the true drift and diffusion functions (black curves). a.u means arbitrary unit.

#### Example 3

In this example we reconstruct a high-resolution ecological dataset. Data is a univariate Cyanobacterial biome measured as phycocyanin concentrations in the Lake Mendota (Carpenter *et al*. 2020). The measurements were taken at minute intervals during the summer thermal stratification of 2011, a period known for common Cyanobacterial blooms (see Figure 3, left panel). For further details on this dataset, refer to references (Arani *et al*. 2021; Magnuson, Carpenter & Stanley 2023).

**Figure 3.**
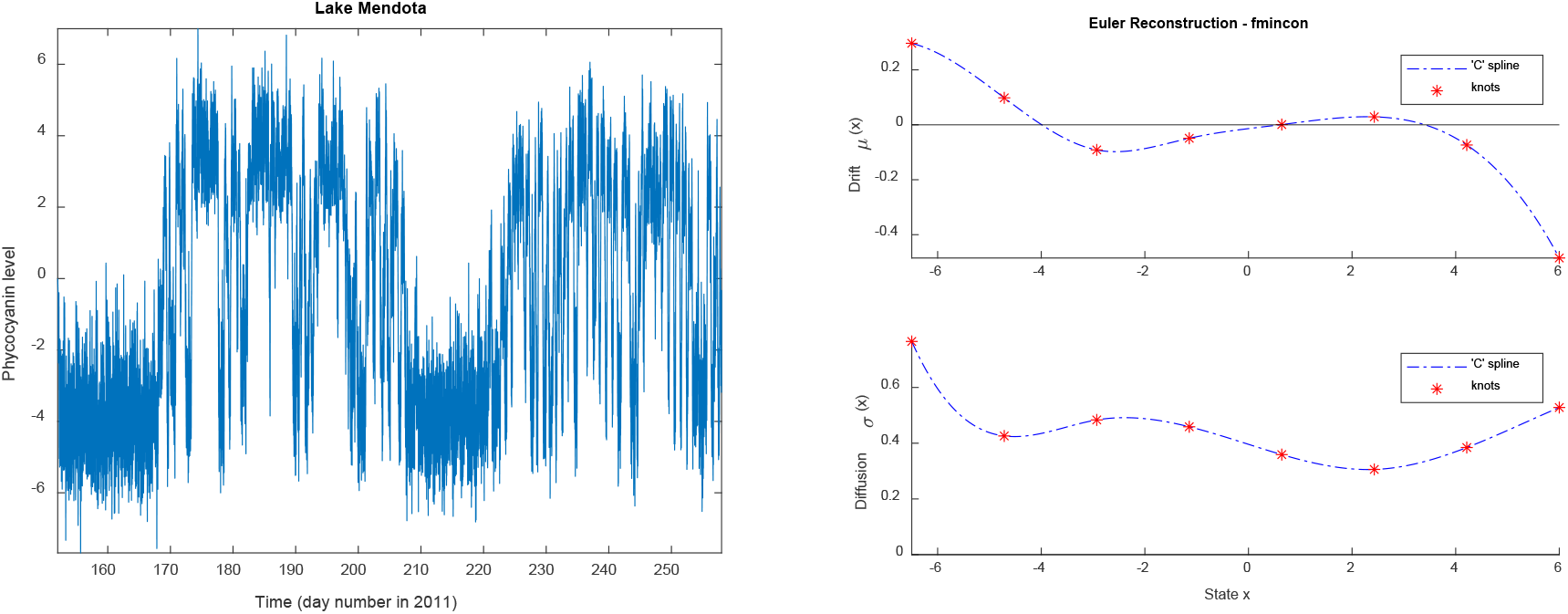
Illustration of spline modeling for a high-resolution ecological dataset. The left panel illustrates a high-resolution cyanobacterial measurement, taken at minute intervals, from lake Mendota. While this dataset does not meet one the data requirements (i.e., Markov property), a rarified sample of this dataset, including every third data point, satisfies this requirement. In the right panel, cubic spline modeling was applied to this rarified sample to estimate the drift and diffusion functions. Data can be found in the reference (Magnuson, Carpenter & Stanley 2023) with the URL https://doi.org/10.6073/pasta/fc8bd96677405945024ad708003be1fc

This dataset does not meet one of the data requirements as it lacks Markov property (indicating high correlations at the measured time scale). However, a rarified sample of this dataset, consisting of every third data point (still maintaining high resolution), does exhibit Markovian behavior. Therefore, we applied Euler reconstruction using cubic splines to the rarified sample (see Figure 3, right panel). The usefulness of spline reconstruction becomes evident here: in real datasets where choosing an appropriate model may be challenging, spline reconstruction proves to be a convenient solution.

### Examples with low-resolution data

#### Example 4

Here, we reconstruct the same dataset in Example 2 but with a resolution being 300 times less. This leads us to a low-resolution dataset. To reconstruct it we follow Hermite reconstruction which has a higher accuracy than Euler reconstruction. Here, we implement spline modeling technique. Notably, we utilize ‘quadratic’ splines instead of the more common cubic splines for Hermite reconstruction, significantly reducing computational time and enhancing the likelihood of successful parameter estimation. The reason has to do with the nature of Hermite reconstruction algorithm (detailed in Appendices F, G, and particularly H). Despite dataset being low-resolution, Hermite reconstruction yields remarkably accurate results in capturing the underlying data-generating system (Figure 4, right panel), outperforming Euler reconstruction (Figure 4, left panel).

**Figure 4.**
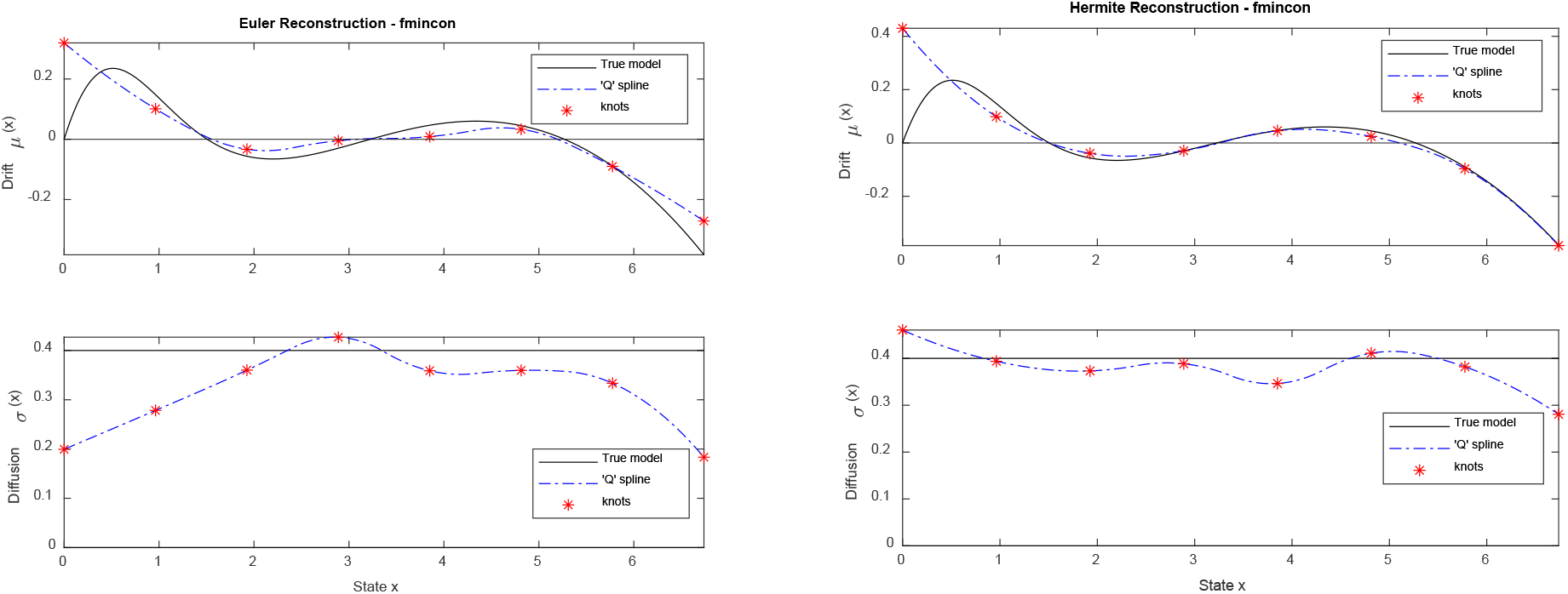
Illustration of Hermite and Euler reconstructions for a low-resolution dataset generated from a nonlinear model. The left panel illustrates Euler estimated quadratic spline models (dot-dashed blue curves) for both the drift and diffusion functions using 8 regularly spaced knots over the state space alongside the true drift and diffusion functions (black curves). The right panel illustrates Hermite estimated quadratic spline models (dot-dashed blue curves) for both the drift and diffusion functions using 8 regularly spaced knots over the state space alongside the true drift and diffusion functions (black curves).

#### Example 5

In this example, a low-resolution univariate climate dataset is reconstructed. The dataset, a *δ*^18^O record from the North Greenland Ice Core Project (NGRIP) (2004), serves as a proxy for the temperature of the northern hemisphere, spanning the last 120 thousand years with a resolution of 20 years. However, this dataset fails to meet two key data requirements outlined in Step2. Initially, the dataset exhibits non-stationarity, although it stabilizes within the period from 70 to 20 thousand years before the present (see Figure 5, top panel). During this epoch, the northern hemisphere climate witnessed alternating colder (stadial) and warmer (interstadial) states, attributed to Dansgaard–Oeschger (DO) events (Dansgaard *et al*. 1993). Within the specified time frame, the majority of DO events, from DO2 to DO18 out of a total of 25 DO events, occurred (2004). Secondly, the dataset lacks the Markov property, yet a rarified sample, with every other point demonstrates Markovian behavior approximately (refer to Table 1 in section 6.3 of the tutorial for further details). Given its low resolution, Hermite reconstruction is deemed suitable. Here, we employ quadratic spline modeling to reconstruct the dataset. Figure 5 illustrates the outcomes of Euler and Hermite reconstructions. Additionally, we introduce a significant and informative quantity known as ‘*effective potential*’ (Arani *et al*. 2021). Unlike deterministic systems where the location of equilibria can be identified using the drift function this is not the case with stochastic systems. For such systems effective potential should be used which is a quantity that incorporates information from both drift and diffusion functions. It is particularly useful for identifying alternative stable states, as is evident in this climate dataset. The minima of effective potential indicate the location of alternative stable states of stadial and interstadial states (solid dots in Figure 6, bottom right panel) which are separated by a repellor in between (open circle in Figure 6, bottom right panel).

**Figure 5.**
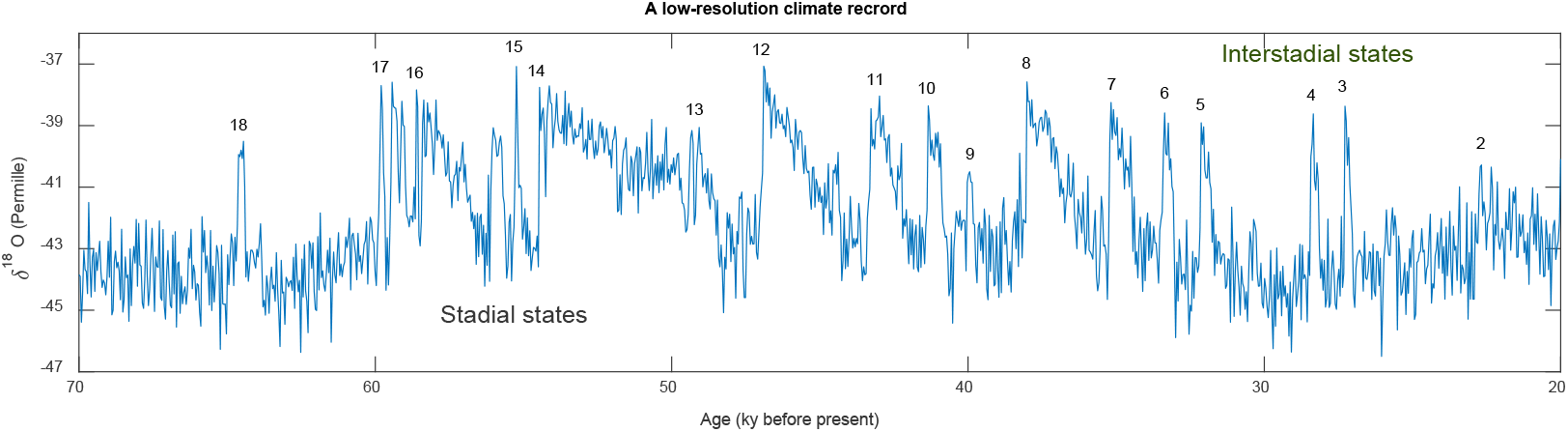
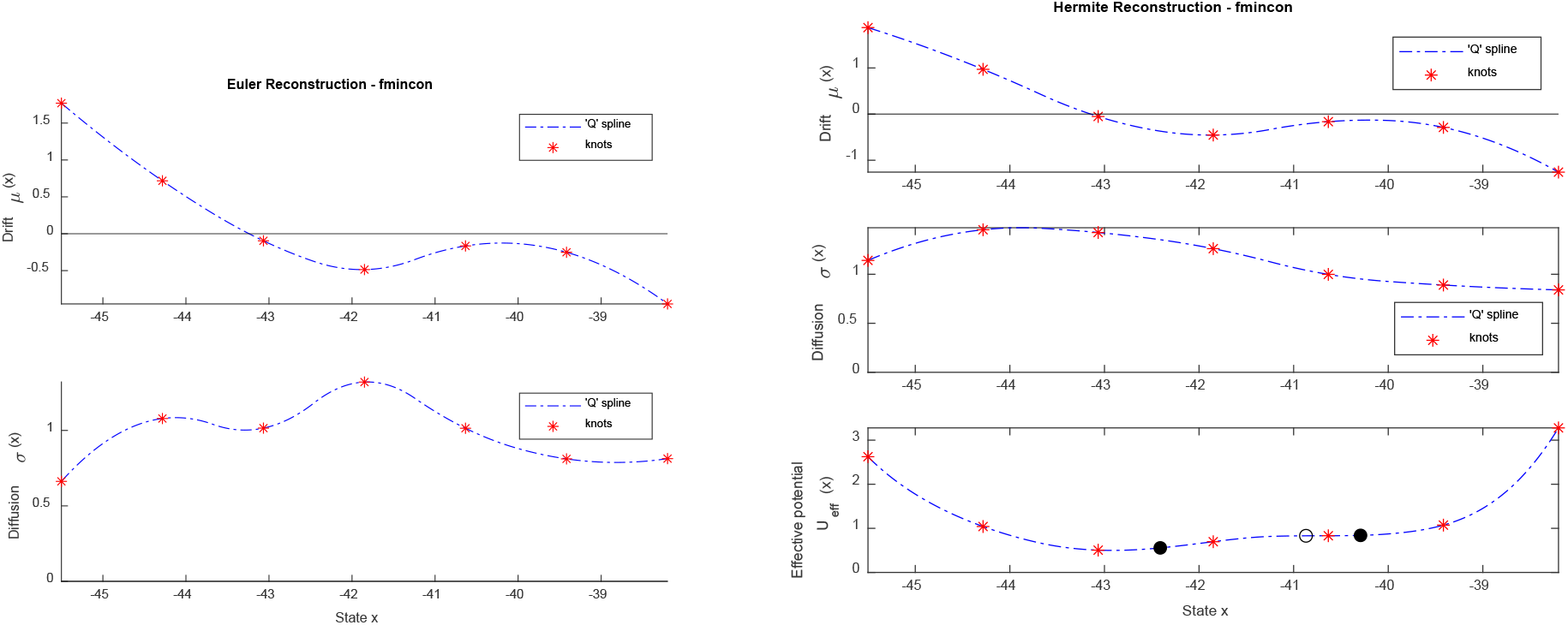
Illustration of Hermite and Euler reconstructions for a low-resolution climate dataset. The top panel illustrates a *δ*^18^O climate record extending from 70 to 20 thousand years before the present time from NGRIP. This is used as a proxy for the temperature of the northern hemisphere which shows that the northern hemisohere climate alternated between cold stadial and warmer interstadial alternative climate states. In this time period majority of Dansgaard-Oescher events occurred (see the labels 2 to 18). The description for bottom left and right panels are similar to that in Figure 5. However, in the bottom right panel the effective potential is also depicted. Effective potential is useful to see whether there are alternative stable states in the dataset which is the case in this dataset (the solid dots represent alternative climate states of stadial and interstadial states separated by the open circle in between).

## Discussion

By delineating a systematic approach encompassing data preparation, model selection, and reconstruction techniques, we furnish researchers with tools and a practical guide for analyzing diverse datasets. The methods and examples presented in this paper furnish valuable insights into the reconstruction of Langevin systems from datasets of varying resolutions. Our discussion of data requirements is fundamental for reconstructing real datasets. We acknowledge the challenges posed by datasets that do not meet these requirements and propose strategies such as data division or rarified sampling to address these limitations. The choice between parametric and spline modeling depends on the nature of the dataset and the researcher’s familiarity with the underlying dynamics, including prior empirical knowledge. While parametric models offer interpretability with respect to model form, spline models provide flexibility in capturing unknown model nonlinearities, rendering them particularly suitable when the true model is uncertain. In general, we recommend to use spline modeling. Furthermore, without incorporation of splines, addressing Hermite reconstruction for low-resolution data pose considerable challenges.

The examples presented illustrate the application of our methodology to diverse datasets, spanning from simulated data to ecological data and climate records. These examples underscore the efficacy of both Euler and Hermite reconstruction techniques, demonstrating their utility across different resolutions and system complexities. Remarkably, Hermite reconstruction proves to be particularly valuable for low-resolution datasets, offering higher accuracy compared to Euler reconstruction. We believe our ‘MATLAB reconstruction package’ together with a step-by-step and user-friendly tutorial is a highly valuable tool for ecologists and life scientists with little affinity for mathematical and statistical modeling.

Overall, our approach furnishes a systematic framework for reconstructing complex systems from observational data. While the examples provided demonstrate the efficacy of our methodology, further research is warranted to explore its applicability to other domains and datasets, especially those generated by more complex processes than Langevin models, such as diffusion-jump models or models driven by Lévy noise. Additionally, ongoing efforts to enhance computational efficiency and address computational challenges associated with multivariate Hermite reconstruction promise to advance the field further.

## Acknowledgements

Lake Mendota data were provided by the North Temperate Lakes Long-Term Ecological Research program from the National Science Foundation under Cooperative Agreement #DEB-2025982.

## Supplementary material for

### A brief on the appendices

**Appendix A** discusses the concept of the ‘relaxation’ time scale, a valuable metric for approximating data resolution before fitting a Langevin model. This metric aids in categorizing data into ’low’ and ‘high’ resolutions, providing essential insights into the feasibility of data reconstruction and the choice of suitable reconstruction algorithms.

**Appendix B** delves into two additional data requirements: data stationarity and data Marcovicity, offering guidance on addressing violations of these prerequisites.

**Appendix C** provides a concise overview of applying maximum likelihood estimation (MLE) for model parameter estimation within Langevin systems.

**Appendix D** outlines a quasi-MLE approach we term the ‘Euler reconstruction’ cautioned against for datasets with low resolution.

**Appendix E** introduces a much more accurate but also more expensive quasi-MLE method pioneered by Aït-Sahalia, we term ‘Hermite reconstruction’ recommended for datasets with lower resolutions.

**Appendix F** presents a refined version of Aït-Sahalia’s approach in Appendix E, enhancing its applicability to general diffusion models at a slight computational cost.

**Appendix G** highlights the advantages of ‘spline’ modeling over traditional ‘parametric’ models, emphasizing superior speed, accuracy, and convenience in using them.

**Appendix H** elaborates on the optimization process for MLE for both Euler and Hermite reconstructions.

For a swift analysis of univariate datasets, we recommend starting with Appendices A and B for foundational insights. Following this, running our MATLAB code ‘AllFigures.m’ will provide in-depth analyses of the five examples outlined in the main text, accompanied by detailed commentary and explanations. If you have the luxury of time and a desire to delve deeper into our package, we encourage you to explore our tutorial. By executing the code lines provided, you can explore numerous additional examples, enriching your understanding of our methodology and its applications. Please find all the codes and the package in our GitHub repository in the following link:

https://github.com/mshoja/MATLAB-reconstruction-package

## Appendix A. Investigating the relaxation time scale of data in advance

It is important to have some knowledge or at least a feeling about the level of data resolution before embarking on the analysis. This can roughly let us know to which regime (possible or impossible to reconstruct. See the main text for the details) our dataset belongs to. And if the answer is yes (i.e., dataset resolution let us to fit a diffusion model), it also helps to find to which resolution category (low, medium, high) our dataset belongs to. In the univariate case, data resolution can be assessed by investigating the autocorrelation of data and then estimating the relaxation (or correlation) time of data: one fits the exponential exp(−*ct*) to some ‘first lags’ of the data autocorrelation function, estimate *c* and finally the relaxation time is *τ_R_* = 1/*c*.

If the data sampling time Δ is much smaller than the relaxation time(s) *τ_R_* (the regime Δ≪ *τ_R_*), then our dataset has a high-resolution and we can safely use simple reconstruction schemes like Langevin approach (see the references (Siegert, Friedrich & Peinke 1998; Rinn *et al*. 2016) for a detailed explanation of this reconstruction scheme but refer to (Arani *et al*. 2021) for a brief overview) or Euler scheme (see Appendix D). When the sampling time is of the same order of magnitude with relation time (the regime Δ≲ *τ_R_* ) there is still hope to infer the system from the data. However, when the sampling time exceeds the relaxation time (the regime Δ> *τ_R_*), it signals that there is very little or no hope to infer the underlying system properly, and all reconstruction techniques might fail. Knowing the relaxation time prior to performing reconstruction is important to select the right reconstruction algorithm. In the first regime, we can safely perform a Euler or Langevin reconstruction, as mentioned. In the second regime, we have to use a more accurate reconstruction procedure like the Hermite reconstruction (Appendices E and F) and use a rather large value of the parameter K (this is a key parameter in Hermite reconstruction). Furthermore, if Δ is between the first and second regime (i.e., data resolution is medium), then we can choose a smaller value of K. Since, Hermite reconstruction with a large K can be time-consuming, having prior knowledge of the data resolution is helpful for selecting the right reconstruction algorithm with proper parameters to strike a balance between accuracy and speed.

## Appendix B. Two more data requirements prior to performing reconstruction

Data should be stationary at least in a weak sense. Loosely speaking, stationarity implies that that the statistical properties of the system (and hence the dataset at hand) should remain constant over time. Weak stationarity over a fixed time-window requires that the mean and variance of data to remain unchanged, and the autocorrelation function depends only on the time lag rather than the initial and final times within the window. However, even if stationarity is violated across the entire dataset, one can consider shorter (overlapping) time windows over which data remain stationary. Reconstruction can then be performed on each window separately, and the results can be interpolated between the windows. Stationarity of the data can be tested using the augmented Dickey-Fuller test (Dickey & Fuller 1979), and this was done for our real datasets (simulated data are, indeed, stationary).

Another requirement is that the noise source in a Langevin model should be uncorrelated or white. This implies that the dataset under study should be Markovian, meaning that the future state should depend only on the current state, independent of past histories. However, in this paper, using Hermite reconstruction, we can relax this assumption: if our main dataset is not Markovian, we can find a coarser time scale called the Markov-Einstein (ME) time scale (Friedrich *et al*. 2011), where Markovicity holds, and perform reconstruction on a sample of data whose time scale is the ME time scale or even coarser.

## Appendix C. Maximum likelihood estimation (MLE) of Langevin models

Consider a Langevin model (the discussion here works for any stochastic Markovian system)

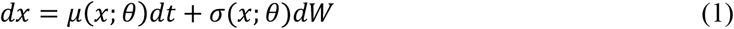

Where *θ* is the vector of unknown parameters. Assume that we have an observation {*X*_0_, *X*_1_, …, *X_N_* } of this model at discrete times *t* = 0,1, …, *N* for a particular set of parameter values which we aim to estimate and let Δ be the sampling time. Assume also that *P_X_*(Δ, *x*|*x*_0_;*θ*) represents the conditional density of *x* ≔ *X*_*t*+Δ_ given *x*_0_ ≔ *X_t_*. The likelihood function for this set of observations is the multi-point density *P_X_*(*X*_0_, *X*_1_, …, *X*;*θ*). Due to the Markovian nature of the Langevin model (1) and using the Bayes rule in probability theory we can greatly simplify this multi-point density by the products of the conditional densities as below

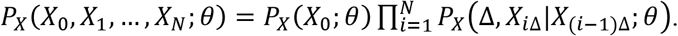

Based on the principle of MLE the true parameters of the Langevin model (1) correspond with the global maximum of the likelihood function. It is more convenient to work with log-likelihood function

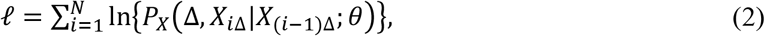

where the first unconditional term in (*P_X_*(*X*_0_)) is ignored as it has a negligible effect when *N* ≫ 1 (see, for instance, the first paragraph in (Aït-Sahalia 2002b)). In this work, we actually consider the negative log-likelihood function −*ℓ* and consider solving a minimization problem, instead. Only for a limited number of Langevin models we know the analytical form of the conditional density in (2). Therefore, our focus lies on quasi-MLE procedures, where we employ approximations of the conditional density in (2) in Appendices D, E, and F.

## Appendix D. Euler reconstruction

The Euler reconstruction corresponds with a maximum likelihood estimation of the Langevin model (1) when the sampling time is infinitesimally small, i.e., Δ→ 0. Under this limiting case we can replace the Langevin model (1) with a difference equation based on the Euler-Maruyama discretization

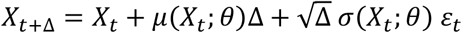

Where ε*_t_* follows a standard normal distribution. As a result, the conditional density *P_X_*(Δ, *x*|*x*_0_;*θ*) can be approximated by a normal distribution with mean *x*_0_ + *μ*(*x*_0_;*θ*)Δ and variance Δ*σ*^2^(*X*_0_;*θ*), that is

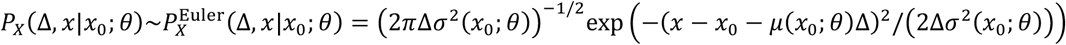

To construct a quasi-MLE procedure one replaces the conditional density in (2) by the approximation 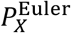. To proceed with parameter estimation this quasi-MLE should be solved. The big advantage of using the Euler approach is due to the fact that its implementation is very fast. Fortunately, based on our experience, this approach also shows a satisfactory performance for datasets with medium resolution.

## Appendix E. Hermite reconstruction

Here, we present an advanced reconstruction procedure developed by Aït-Sahalia (Aït-Sahalia 2002a). For the sake of clarity and convenience we keep exactly the same notation as Aït-Sahalia. As discussed in Appendix C, the process of a quasi-MLE requires the estimation of conditional density *P_X_*(Δ, *x*|*x*_0_;*θ*). The approach of Aït-Sahalia is more accurate than the Euler approach in Appendix D and is based on establishing a convergent Hermite series expansion for the conditional density *P_X_*(Δ, *x*|*x*_0_;*θ*) and hence a convergent expansion for the log-likelihood function *ℓ* in (2) using Hermite polynomials. Therefore, we call this approach ‘Hermite reconstruction’. This is achieved through two transformations, *X* → *Y* and *Y* → *Z*, where the conditional distributions in the transformed space *Z* will be as close as to the standard normal distribution. Aït-Sahalia then constructs a convergent Hermite series expansion for the transformed variable *Z* and finally back-transforms to the original variable *X*. The first transformation *X* → *Y*, also known as the Lamperti transform, is as follows

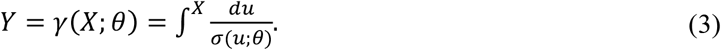

By the Itô’s lemma, the corresponding Langevin equation (1) for the transformed variable *Y* has a unit diffusion, i.e., *dY* = *μ_Y_*(*Y*;*θ*)*dt* + *dW* and

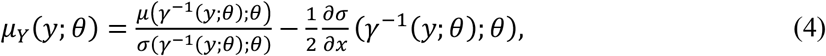

where *γ*^−1^ is the inverse function of *γ*. The conditional density of the variable *Y* is now closer to normal distribution due to its unit diffusion. Since the distribution of *Y* can get peaked when the sampling time Δ is very small Aït-Sahalia performs a second transformation *Y* → *Z* to overcome this problem. Stated other way, he standardizes *Y* as

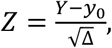

where *y*_0_ is the corresponding value of *x*_0_ following the transformation *Y*. Aït-Sahalia shows that *Z* is close enough to standard normal distribution and then expands the density of *Z, P_Z_*, about standard normal distribution *ϕ*(*z*). Therefore, the conditional density *P_Z_*(Δ, *z*|*y*_0_;*θ*) of *Z* can be expanded about the standard normal distribution, by the Hermite polynomials up to the *J*^th^ term as below

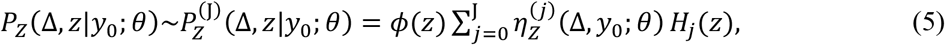

where *H_j_* (*z*) are the classical Hermite polynomials being defined as 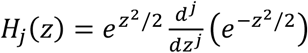 *j* ≥ 0 so the first few Hermite polynomials read *H*_0_(*z*) = 1, *H*_1_(*z*) = −*z, H*_2_(*z*) = *z*^2^ − 1, *H*_3_(*z*) = −*z*^3^ + 3*z*, …. Since *H*_0_ (*z*) = 1 we have that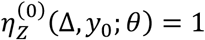. For *j* > 1, the coefficients 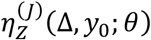 can be found by the orthogonal properties of Hermite polynomials as bellow

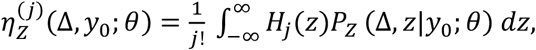

which is a conditional expectation. Each of the Hermite coefficients 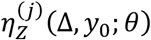 can be approximated using Taylor expansion in Δ up to the K^th^ term as bellow (see also formula 4.21 in (Jeisman 2006))

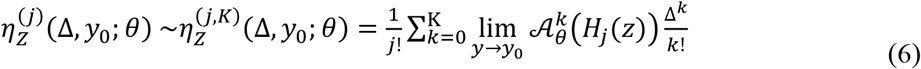

Where 𝒜*_θ_* is the ‘*infinitesimal generator*’ of the process *Y* (for an overview on the infinitesimal generator and its properties see, for instance, page 18 in (Jeisman 2006)) which is defined as

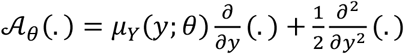

Note that the notation 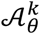 in (6) is the *k^th^* self-composition of the operator 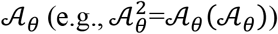. Finally, we arrive at the following approximation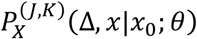 for our quantity of interest, i.e., *P* (Δ, *x*|*x*_0_;*θ*), using the Jacobian formula and by specifying how many Hermite terms (J) in (5) and how many temporal terms (K) in (6) we wish to include

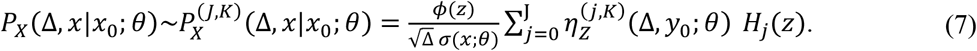

As Aït-Sahalia argues (and we experienced) the expansion in (5) can be well approximated by including at most the first three terms (so, *J* = 1,2,3 are often enough). It is important to note that using (6) all coefficients 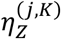 of the Hermite expansion of conditional density in (7) can be expressed by *μ_Y_* and its higher derivatives at *y*_0_ up to order 2K-2, i.e., *μ_Y_*(*y*_0_), *μ_Y_* ′ (*y*_0_), …, *μ_Y_*^[2K−2]^(*y*_0_). We will discuss on this extremely important point in more details in Appendix G.

## Appendix F. A refinement to Hermite reconstruction

As discussed in Appendix E, the approach by Aït-Sahalia necessitates the availability of the function *γ* in (3), which maps the process *X* to the process *Y*, in an analytical form. Additionally, even if *γ* is analytically available, the method requires the inverse function of *γ*,i.e., *γ*^−1^, to be known analytically as well. Consequently, this procedure imposes no restrictions on the drift function, and possible restrictions are related to the diffusion function. While this might not pose a significant limitation for most stochastic models encountered in ecology and biology, including additive Langevin models, there may be cases where such analytical requirements are not feasible (see (Bakshi & Ju 2005)). Moreover, even for models without such restrictions, it remains advantageous to consider the methodology presented in this appendix. This is because the approach here differes slightly from that of Aït-Sahalia, offering enhanced accuracy albeit with slightly longer computational time. Here, we provide a brief overview of the method, omitting detailed explanations. The refinement keeps the transformation *X* → *Y*, but considers the following standardization of *Y* by its true mean and variance, in contrast to unlike Aït-Sahalia’s approach, which he refers to as ‘pseudo-normalization’(Aït-Sahalia 2002a))

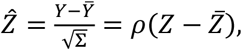

where 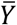 and Σ are, respectively, the true mean and variance of *Y*, 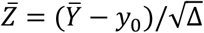 is the mean of 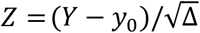 and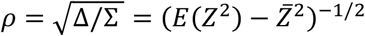. The variable *Z* can be rewritten in the following form so that it would be a direct function of *X* and *x*_0_ rather than *Y* and *y*_0_

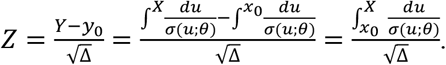

Note that in this approach there is no need to know the integral 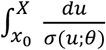 analytically and only a numerical approximation is enough. The way this refinement standardizes *Y* and rewrites *Z* allows to do all the calculations directly in terms of the original drift function *μ*(*x*) and diffusion function *σ*(*x*) and avoids expressing the results in terms of *μ_Y_*(*y*) in (4), i.e., the drift function of the process *Y*. A Hermite expansion approximation up to the order J for the conditional density *P_X_*(Δ, *x*|*x*_0_;*θ*), using the Jacobian formula applied to the density of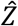, reads

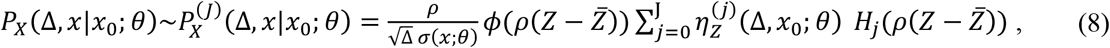

in which the Hermite coefficients are obtained as

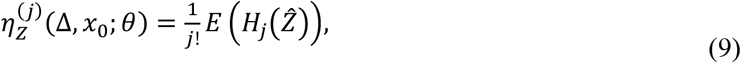

where *H_j_* are classical Hermite polynomials of order *j* ≥ 0 (note that in (8) the first and second coefficients 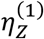 and 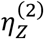 can be shown to be 0 (Bakshi & Ju 2005). Therefore, based on this algorithm *j* actually starts from 3). Stated other way, the Hermite expansion coefficients *η*_*j*_(Δ, *x*_0_;*θ*) in (9) are expressed in terms of moments of the powers of 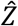, where the later can be expressed by the moments of the powers of *Z*, i.e., *E*(*Z*^*i*^,), *i* ≥ 0. *E*(*Z*^*i*^,) can be approximated by a Taylor expansion in Δ up to order *K* as

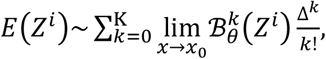

where ℬ*_θ_* is the ‘*infinitesimal generator’* of the process *X* which is defined as

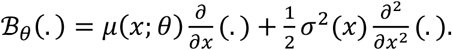

Clearly, this leads us to an approximation in terms of the parameters J and K, say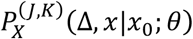, for the conditional density of *X* in (8) where the Hermite coefficients 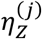are approximated by 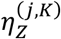. It is important to note that all Hermite expansion coefficients 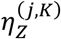 can be expressed by *μ*(*x*) and its higher order derivatives at *x*_0_ up to order 2K-2 and by *σ*(*x*) and its higher derivatives at *x*_0_ up to order 2K-1. We will discuss on this extremely important point in more details in Appendix G.

## Appendix G. Several reasons for the superiority of spline modeling in Hermite reconstruction

In Appendix E we argued that using the approach by Aït-Sahalia Hermite expansion coefficients can be expressed by the drift function of the process *Y*, i.e., *μ_Y_*, and its higher order derivatives up to order 2K-2, i.e., *μ_Y_, μ_Y_* ′, …, *μ_Y_*^[2K−2]^. Likewise, we also argued in Appendix F that using the refined approach the corresponding Hermite expansion coefficients can be expressed by the drift and diffusion functions of the process *X* up to orders 2K-2 and 2K-1, respectively, i.e., *μ, μ* ′, …, *μ*^[2*KK*−2]^, *σ, σ* ′, …, *σ*^[2*KK*−1]^. Therefore, using both algorithms, as the differentiation order increases, it generally leads to significant computational complexities, especially as K increases. The computational complexities also increase as the number of Hermite terms (J) in the expansion of conditional density increases. However, in practice, the first or first two Hermite terms are sufficient in both algorithms (Aït-Sahalia 2002b) (i.e., in the first algorithm we at most need J=1,2 and in the second one we at most need J=3,4). Consider, for instance, a stochastic version of the overgrazed model of May (May 1977) with drift part 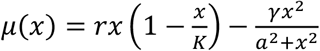. As the differentiation order of *μ*(*x*) increases, the algebraic expressions become longer, imposing a substantial computational burden. In contrast, higher order derivatives of a cubic polynomial (which approximately fits the May model) result in shorter expressions that eventually vanish at order = 4. This suggest using polynomial forms for *μ_Y_*(*y*) in the first algorithm and polynomial forms for *μ*(*x*) and *σ*(*x*) in the second algorithm.

While polynomials are convenient candidate models, they may not be ideal due to the need for higher-order polynomials to adequately represent the nonlinearities of complex models. Instead, piecewise polynomials, or splines, are ideal candidate models due to their flexibility and simplicity. Splines offer great accuracy, comfort, and speed in the reconstruction process. They are ‘flexible’ enough to capture unknown linearities in the data-generating process and have a ‘simple’ polynomial form in their building blocks, which reduces computational burden significantly. Since cubic splines typically suffice in practice, they are more attractive than polynomials from a computational standpoint. If { *x*_1_, *x*_2_, …, *x*_*M*_ } denotes a ‘knot sequence’ across the state space then the model parameters in spline reconstruction are the unknown values of drift and diffusion functions at these knots. It worth mentioning that the reconstruction approach using spline forms is parametric. However, the modeling feels ‘non-parametric’ as if there is no explicit model involved. Therefore, we call it ‘spline modeling’ to distinguish it from ‘typical’ parametric models.

Here, we highlight additional reasons for the superiority of spline modeling beyond speed. The second reason for the superiority of splines is that splines are linear functions in terms of parameters. This significantly enhance the optimization process as it often leads to an optimization problem with one or very few local minima. Third, in general we have no idea how to choose a proper model to try, unless there is robust empirical or theoretical justification. Even in cases where a parametric model is preferred, it is recommended to consider fitting a spline model to the data first to gain insights into a proper parametric model form to try. The fourth reason is that in the process of estimating model parameters using MLE, the optimization algorithm needs to search within a sufficiently large parameter space containing the true but yet unknown parameters. However, it is generally challenging to define a proper bounded parameter region around the true solution when considering a typical parametric model. In contrast, spline modeling offers a convenient way to establish a proper bounded parameter space for the algorithm to search within. The reason is simple: in spline modeling, our parameters hold a special meaning as they represent the values of drift and diffusion functions at knots. The fifth reason is that, when using splines, our models will not exhibit global sensitivity to parameters. A change in the value of a parameter at a single knot will not propagate across the entire knot sequence. Instead, the effect is localized over an evenly spaced knot sequence, which we always choose (De Boor & De Boor 1978). A model with sharp sensitivity to parameters places significant pressure on optimization procedure in terms of both accuracy and speed, potentially leading to failure.

### Quadratic splines: A more efficient approach for Hermite reconstruction than cubic splines

Although the use of cubic spline forms leads to a huge reduction in the computational complexity, we might still need a further reduction of computational burden. Quadratic splines have a slightly less desirable smoothing property in comparison with cubic splines but the use of quadratic splines relative to cubic splines greatly reduces the computations as parameters J and/or K increase. Let’s assume we want to follow the methodology in Appendix E (the same logic works if we follow the methodology in Appendix F). Using cubic spline forms for drift and diffusion functions one does not need to express the Hermite coefficients in terms of higher derivatives of drift and diffusion functions up to orders 2K-2 and 2K-1, respectively. Instead, the Hermite expansion coefficients can be expressed by the derivatives of the drift and diffusion functions up to order 3 only, i.e., *μ, μ* ′, *μ ″, μ ‴, σ, σ ′, σ ″, σ ‴*, and we get rid of higher order derivatives. Clearly, this leads us to a huge simplification of conditional density in (8) especially when J and/or K is big. On the other hand, a further level of simplification is achievable by using quadratic spline forms since we can get rid of third order derivatives and express the Hermite coefficients by *μ, μ ′, μ ″, σ, σ ′, σ ″* only. In additive models (in either case of cubic and quadratic splines) an even higher level of simplicity is attainable since *σ ′*= *σ ″* = 0. Therefore, the reduction of computational burden from cubic splines to quadratic splines will not be felt sharply for additive models. However, in multiplicative models, which we often desire, such a reduction makes a big difference. Because of this huge reduction of computational time, we could often tackle Hermite reconstruction using quadratic splines rather than cubic splines.

### A note on data standardization for spline modeling

When the range of data encompasses big numbers there is a risk for numerical instabilities. In such cases, it is better to standardize the data first, perform the analysis on the standardized data and, at the end back transform the results to the original scale of data. This is especially handy for linear models such as spline models. Consider the standardization of a state variable *x*, i.e., *z* = (*x* − *m*_data_)/*s*_data_ in which *m*_data_ and *s*_data_ are the mean and standard deviation for a dataset of the process *x*. Now, assume that the Langevin model *dz* = *μ_z_*(*z*)*dt* + *σ_z_*(*z*)*dW* describes the dynamics of the transformed process *z*. Then the corresponding Langevin model for the process *x* has the following drift and diffusion functions

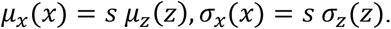

Assume that { *x*_1_, *x*_2_, …, *x*_*M*_ } and {*z*_1_, *z*_2_, …, *z_M_* } are the knot sequences for the processes *x* and *z*. Assume further that the estimated drift and diffusion parameters of a spline model for the corresponding standardized data are *A* = {*μ*_1*z*_, *μ*_2*z*_, …, *μ_Mz_*} and *B* = {*σ*_1*z*_, *σ*_2*z*_ …, *σ_Mz_*}, respectively (these are the estimated values of the drift and diffusion functions of the *zz* process at the knot sequence {*z*_1_, *z*_2_, …, *z_M_* }). Now, to back transform everything to the original scale (i.e., the *x* process) all we need to do is to multiply the elements of *A* and *B* by the data standard deviation *s*_data_ to find the estimated parameters of the process *x* over the knot sequence { *x*_1_, *x*_2_, …, *x*_*M*_ }.

## Appendix H. Gradient descent and grey wolf optimizer algorithms used to solve the MLE

In order to estimate the parameters of the Langevin model (1) using the MLE framework we should find the global minimum of negative log-likelihood function −*ℓ* where *ℓ* is defined in (2) in Appendix C. The approach to tackling the MLE problem significantly differs between Euler and Hermite reconstructions. Euler reconstruction, the objective function is fully defined across the entire parameter space. However, in Hermite reconstruction, the objective function may become undefined for certain parameter values as they deviate from the true global minimum, a discrepancy that becomes more pronounced as data resolution decreases. This is because as the data resolution declines the Hermite expansion of conditional density in (8), or in (7), might converge to a density which may not be entirely positive across the state space (although it always integrates to 1), leading to a partially defined objective function. Typically, within a small enough neighborhood around the optimum parameter values, the Hermite series converges to a positive density. Yet, as this neighborhood expands, the series may not maintain a positive density for some parameter values, which we term ‘illegitimate’ solutions (as opposed to ‘legitimate’ solutions). In practice, the size and geometry of the region where objective values are defined, i.e., the legitimate region, have a complex dependency on factors such as data resolution, model complexity, and the level of approximation adopted in selecting the K and J values in the Hermite reconstruction algorithm. Therefore, in the case of Hermite reconstruction, we face not only a non-smooth objective function but also a partially defined one, where the density of undefined (i.e., illegitimate) objectives increases the further we move from the global minimum. To handle this constraint, we employ a ‘death penalty’ technique, which assigns a very high positive number to illegitimate objective values. Consequently, classical optimization routines based on the gradient descent algorithm cannot straightforwardly solve this complex optimization problem.

To enable the use of gradient descent algorithms in our partially-defined optimization problem, we employ a two-phase algorithm. The initial phase involves addressing the Euler reconstruction to obtain the parameter estimate 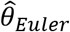. For this optimization, we adopt a strategy of employing multiple starting points. This approach helps us to navigate around the potential issue of being trapped in a local minimum. However, based on our extensive experience, when the model is linear with respect to its parameters, we typically encounter an optimization landscape characterized by a unique local minimum that also serves as the global minimum or relatively few local minima. This observation underscores the practical benefits of working with such linear models, particularly spline models, which tend to simplify the optimization process significantly.

In the second phase, we focus on exploring the vicinity of 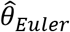, where we anticipate a higher density of legitimate solutions. Our aim here is to identify several legitimate solutions that can serve as a basis for constructing a surrogate model. The feasibility and effectiveness of this surrogate model hinge on the concentration of legitimate solutions near 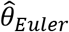. Should we successfully establish a surrogate model, it often becomes possible to identify its global minimum 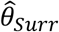, which usually lies in close proximity to the true global minimum. This surrogate solution becomes a valuable starting point for deploying a gradient descent algorithm on the original optimization problem. In instances where a surrogate model proves unattainable, the legitimate solutions already identified are utilized as initial points for a gradient descent algorithm on the original optimization problem. Regardless of the presence of a surrogate model, these starting points, which are hoped to have a high density of legitimate points near them, can pave the way for the application of the gradient descent algorithm. Our aim is to improve 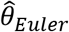 through this process, aspiring to discover a new solution 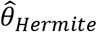, which either improves upon 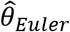 or, in the worst case, matches it. Should the complexity of the chosen model prevent any enhancement of 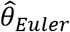, we consider simplifying the model and iterating the process to achieve a superior solution. Our extensive experience, supported by numerous case studies, demonstrates that this algorithm frequently yields superior outcomes when spline modeling is employed, as opposed to traditional parametric models. This is extensively documented through various examples in section 11 of our tutorial.

### Accessing the uncertainty of the parameters

To calculate the variance of the estimated parameter vector, say 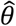, the following Fisher information (FI) matrix is needed

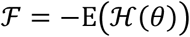

where *ℋ* (*θ*) = ∂^2^*ℓ*/∂*θ*∂*θ^T^* is the Hessian matrix, i.e., the second order partial derivatives of the log-likelihood function with respect to *θ* (in case of minimization, as is our case, −*ℓ* is used as objective function so that the ℱ matrix does not include the negative sign). The ‘observed’ Fisher information approximates the ℱ matrix as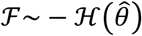. Consequently, the variance-covariance matrix, Σ = ^*ℱ*−1^, i.e., the inverse of the Fisher information matrix, is estimated by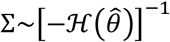. The standard error of the i*^th^* parameter, say 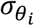, which is the square root of the i*^th^* diagonal element in Σ, is thus estimated from the square root of the i*^th^* diagonal element of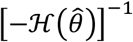. To assess the uncertainties in the case of Hermite reconstruction, extra caution is necessary when estimating the *ℱ* matrix. The crux of challenge lies in ensuring that the objective values used for estimating the *ℱ* matrix are legitimate, not just at the estimated parameters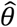, which is known to be legitimate, but also at neighboring parameter values in close proximity to 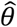, which we are not sure. To tackle this complexity, if an illegitimate objective value is encountered, we apply an infinitesimal perturbation to the parameter in question, ensuring the perturbed parameter yields a legitimate objective.

### An interesting point concerning the uncertainty of drift and diffusion parameters

In our analysis across numerous examples, we have observed a higher accuracy in the estimation of the noise parameters compared to that of drift parameters. This phenomenon is not purely empirical but is supported by theoretical underpinnings. For readers interested in a deeper mathematical explanation, we direct them to references (Sorensen 2007; Tang & Chen 2009; Chang & Chen 2011), where detailed justifications are provided.

## References

(2004) High-resolution record of Northern Hemisphere climate extending into the last interglacial period. Nature, 431, 147–151.

Aït-Sahalia, Y.J.E. (2002) Maximum likelihood estimation of discretely sampled diffusions: a closed-form approximation approach. 70, 223–262.

Arani, B.M., Carpenter, S.R., Lahti, L., Van Nes, E.H. & Scheffer, M. (2021) Exit time as a measure of ecological resilience. Science, 372, eaay4895.

Bakshi, G. & Ju, N.J.T.J.o.B. (2005) A Refinement to Aït-Sahalia’s (2002)”Maximum Likelihood Estimation of Discretely Sampled Diffusions: A Closed-Form Approximation Approach”. 78, 2037–2052.

Bandi, F.M. & Phillips, P.C. (2003) Fully nonparametric estimation of scalar diffusion models. Econometrica, 71, 241–283.

Bolker, B.M., Gardner, B., Maunder, M., Berg, C.W., Brooks, M., Comita, L., Crone, E., Cubaynes, S., Davies, T. & de Valpine, P. (2013) Strategies for fitting nonlinear ecological models in R, AD M odel B uilder, and BUGS. Methods in Ecology and Evolution, 4, 501–512.

Box, G.E. & Cox, D.R. (1964) An analysis of transformations. Journal of the Royal Statistical Society Series B: Statistical Methodology, 26, 211–243.

Carpenter, S.R., Arani, B.M., Hanson, P.C., Scheffer, M., Stanley, E.H. & Van Nes, E. (2020) Stochastic dynamics of Cyanobacteria in long-term high-frequency observations of a eutrophic lake. Limnology and Oceanography Letters, 5, 331–336.

Connell, J.H. & Sousa, W.P. (1983) On the evidence needed to judge ecological stability or persistence. The American Naturalist, 121, 789–824.

Dansgaard, W., Johnsen, S.J., Clausen, H.B., Dahl-Jensen, D., Gundestrup, N.S., Hammer, C.U., Hvidberg, C.S., Steffensen, J.P., Sveinbjörnsdottir, A. & Jouzel, J. (1993) Evidence for general instability of past climate from a 250-kyr ice-core record. Nature, 364, 218–220.

Ditlevsen, P.D. (1999) Observation of α-stable noise induced millennial climate changes from an ice-core record. Geophysical Research Letters, 26, 1441–1444.

Friedrich, R., Peinke, J., Sahimi, M. & Tabar, M.R.R.J.P.R. (2011) Approaching complexity by stochastic methods: From biological systems to turbulence. 506, 87–162.

Hilborn, R. & Mangel, M. (2013) The ecological detective: confronting models with data (MPB-28). Princeton University Press.

Magnuson, J.J., Carpenter, S.R. & Stanley, E.H. (2023) North Temperate Lakes LTER: High Frequency Data: Meteorological, Dissolved Oxygen, Chlorophyll, Phycocyanin-Lake Mendota Buoy 2006-current. 10.6073/pasta/fc8bd96677405945024ad708003be1fc

May, R.M. (1977) Thresholds and breakpoints in ecosystems with a multiplicity of stable states. Nature, 269, 471–477.

Rinn, P., Lind, P.G., Wächter, M. & Peinke, J.J.a.p.a. (2016) The Langevin Approach: An R Package for Modeling Markov Processes.

Scheffer, M. (2009) Critical transitions in nature and society. Princeton University Press.

Scheffer, M., Carpenter, S., Foley, J.A., Folke, C. & Walker, B.J.N. (2001) Catastrophic shifts in ecosystems. 413, 591.

Siegert, S. & Friedrich, R.J.P.R.E. (2001) Modeling of nonlinear Lévy processes by data analysis. 64, 041107.

## References

Aït-Sahalia, Y. (2002a) Maximum likelihood estimation of discretely sampled diffusions: a closed-form approximation approach. Econometrica, 70, 223–262.

Aït-Sahalia, Y.J.E. (2002b) Maximum likelihood estimation of discretely sampled diffusions: a closed-form approximation approach. 70, 223–262.

Arani, B.M., Carpenter, S.R., Lahti, L., Van Nes, E.H. & Scheffer, M.J.S. (2021) Exit time as a measure of ecological resilience. 372, eaay4895.

Chang, J. & Chen, S.X. (2011) On the approximate maximum likelihood estimation for diffusion processes.

De Boor, C. & De Boor, C. (1978) A practical guide to splines. springer-verlag New York.

Dickey, D.A. & Fuller, W.A.J.J.o.t.A.s.a. (1979) Distribution of the estimators for autoregressive time series with a unit root. 74, 427–431.

Jeisman, J.I. (2006) Estimation of the parameters of stochastic differential equations. Queensland University of Technology.

Lehle, B.J.J.o.S.P. (2013) Stochastic time series with strong, correlated measurement noise: Markov analysis in n dimensions. 152, 1145–1169.

Sorensen, M. (2007) Efficient estimation for ergodic diffusions sampled at high frequency. CREATES Research Paper.

Tabar, R. (2019) Analysis and data-based reconstruction of complex nonlinear dynamical systems. Springer.

Tang, C.Y. & Chen, S.X. (2009) Parameter estimation and bias correction for diffusion processes. Journal of Econometrics, 149, 65–81.

